# *2b* or not *2b*: Experimental evolution of functional exogenous sequences in a plant RNA virus

**DOI:** 10.1101/079970

**Authors:** Anouk Willemsen, Mark P. Zwart, Silvia Ambrós, José L. Carrasco, Santiago F. Elena

**Affiliations:** Instituto de Biología Molecular y Celular de Plantas (IBMCP), Consejo Superior de Investigaciones Científicas-Universidad Politécnica de Valencia, Campus UPV CPI 8E, Ingeniero Fausto Elio s/n, 46022 València, Spain; Present address: MIVEGEC (UMR CNRS 5290, IRD 224, UM), National Center for Scientific Research (CNRS), 911 Avenue Agropolis, BP 64501, 34394 Montpellier, Cedex 5, France; Present address: Institute of Theoretical Physics, University of Cologne, Zülpicher Straße 77, 50937 Cologne, Germany; Instituto de Biología Integrativa y de Sistemas (I^2^SysBio), Consejo Superior de Investigaciones Científicas-Universitat de València, Parc Científic de la Universitat de València, Catedrático Agustín Escardino 9, 46980 Paterna, València, Spain; The Santa Fe Institute, 1399 Hyde Park Road, Santa Fe, NM 87501, USA

**Keywords:** Horizontal gene transfer, Virus evolution, Genome evolution, Experimental evolution

## Abstract

Horizontal gene transfer (HGT) is pervasive in viruses, and thought to be a key mechanism in their evolution. On the other hand, strong selective constraints against increasing genome size are an impediment for HGT, rapidly purging horizontally transferred sequences and thereby potentially hindering evolutionary innovation. Here we explore experimentally the evolutionary fate of viruses with simulated HGT events, using the plant RNA virus *Tobacco etch virus* (TEV), by separately introducing two functional, exogenous sequences to its genome. One of the events simulates the acquisition of a new function though HGT of a conserved AlkB domain, responsible for the repair of alkylation or methylation damage in many organisms. The other event simulates the acquisition of a sequence that duplicates an existing function, through HGT of the 2b RNA silencing suppressor from *Cucumber mosaic virus* (CMV). We then evolved these two viruses, tracked the maintenance of the horizontally transferred sequences over time, and for the final virus populations, sequenced their genome and measured viral fitness. We found that the AlkB domain was rapidly purged from the TEV genome, restoring fitness to wild-type levels. Conversely, the *2b* gene was stably maintained and did not have a major impact on viral fitness. Moreover, we found that *2b* is functional in TEV, as it provides a replicative advantage when the RNA silencing suppression domain of HC-Pro is mutated. These observations suggest a potentially interesting role for HGT of short functional sequences in ameliorating evolutionary constraints on viruses, through the duplication of functions.

## Introduction

Viruses play an important evolutionary role as vectors for horizontal gene transfer (HGT) in the genomes of theirs hosts. For example, integrated virus genomes are commonly found in prokaryotes, as prophages can constitute 10-20% of bacterial genomes (Canchaya et al. 2003; Casjens 2003), which serve as vectors for HGT between bacteria and contribute to the genetic variability in several bacterial species (Muniesa et al. 2013; Penadés et al. 2015). Among eukaryotes, HGT mediated by dsRNA viruses is widespread, and may play significant roles in their evolution (Liu et al. 2010). Baculoviruses, dsDNA viruses, have been shown to play a role in HGT of transposons between their insect hosts (Gilbert et al. 2014). In addition to the transfer from viruses to host cells, HGT also occurs commonly from host cells to viruses and from viruses to viruses. For example, phylogenetic analysis suggest that many Mimivirus genes were acquired by HGT from the host organism or from bacteria that parasitize the same host (Moreira & Brochier-Armanet 2008). Another example of HGT from host to virus is illustrated by plant closteroviruses, that encode homologs of Hsp70 (heat shock protein, 70 kDa) (Dolja et al. 1994, 2006). In general, viruses do not encode Hsp70s but recruit host Hsp70s to aid virion assembly or genome replication (Cripe et al. 1995; Kelley 1998). In the case of closteroviruses, the *hsp70* gene was probably acquired by a common ancestor through recombination with a host mRNA coding for Hsp70 (Dolja et al. 1994, 2006).

Even though HGT is a widespread key mechanism in virus evolution, strong selection against increasing genome sizes in viruses is an impediment to HGT (Holmes 2003; Zwart et al. 2014; Willemsen et al. 2016b). An exogenous sequence that is transferred must be accommodated into the genome of the recipient virus. We postulate that the accommodation of gene sequences could entail the regulation of expression levels, fine-tuning of interactions with other viral or host proteins, and the optimization of codon usage. However, exogenous sequences in a viral genome will generally have an appreciable fitness cost, typically resulting in the rapid loss of these exogenous sequences (Dolja et al. 1993; Zwart et al. 2014). When a new sequence which may potentially be beneficial for the virus is incorporated in the genome, a race against time begins: will the new sequence be accommodated or will it be purged? The high instability of viral genomes may therefore hinder evolutionary innovation, by shortsightedly purging elements that might have been beneficial in the longer term. Previous studies that highlighted instability of exogenous sequences used marker genes, as these sequences can reasonably be assumed to be unable to acquire functionality.

In this study we experimentally explore the evolutionary fate of exogenous genes introduced into the genome of *Tobacco etch virus* (TEV; genus *Potyirus*; family Potyviridae), by considering sequences with functions that could potentially be beneficial to the virus. We have simulated two possible HGT events. The first event represents the acquisition of a heterologous gene that duplicates a function that was already present in TEV, thus generating functional redundancy. The second event adds a novel function to the viral genome.

The first HGT event duplicates the RNA-silencing suppression activity. Plant RNA viruses counteract the host plant’s immune responses by encoding viral suppressors of RNA silencing (VSRs). Most plant RNA viruses encode at least one VSR, but others like the closteroviruses, criniviruses and begomoviruses encode more than one (Ding & Voinnet 2007; Díaz-Pendón & Ding 2008). In the case of TEV, the multi-functional HC-Pro protein is the only VSR. HC-Pro can be divided into three regions: (*i*) an N-terminal region associated with aphid transmission, (*ii*) a C-terminal region associated with proteinase and VSR activities (Varrelmann et al. 2007), and (*iii*) a central region implicated in many functions, including VSR activity (Plisson et al. 2003). The silencing suppressor activity of HC-Pro is not always strong enough to nullify silencing effects on virus replication (Gammelgard et al. 2007). Moreover, the VSR activity of this multifunctional protein may be subject to pleiotropic constraints, which arise due to its other functions. We have chosen to insert the *2b* gene from *Cucumber mosaic virus* (CMV; genus *Cucumovirus*, family *Bromoviridae*) into TEV genome. The multi-functional 2b protein is implicated in polyprotein cleavage, virulence, viral movement, aphid-borne transmission and suppression of RNA silencing (Ding et al. 1995, 1996; Li et al. 1999; Guo & Ding 2002; Shi et al. 2003). The addition of a second silencing suppressor to the TEV genome could have both immediate benefits for replication, as well as second-order evolutionary benefits. Immediate benefits could arise simply due to stronger silencing-suppression activity based upon complementary molecular mechanisms. Second-order benefits could occur during evolution, because pleiotropic constraints on HC-Pro’s other functions have been ameliorated by the introduction of the *2b* gene and these functions can therefore be improved.

The second genome reorganization event represents the acquisition of a function that was not originally present in TEV. We have chosen to add an *AlkB* domain. The AlkB protein is responsible for the repair of alkylation damage in DNA and RNA bases. Alkylating agents that produce this damage can be present in the environment and inside cells (Sedgwick et al. 2007). Homologues of *AlkB* are widespread in eukaryotes and bacteria (Aravind & Koonin 2001; Kurowski et al. 2003), as well as in plant RNA viruses (Aravind & Koonin 2001). Although the overall sequence similarity among the *AlkB* family is low, a conserved domain exists characterized as the 2OG-Fe(II) oxygenase superfamily (Aravind & Koonin 2001; Finn et al. 2014). The 2-oxoglutarate (2OG) and Fe(II)-dependent oxygenases are a class of enzymes that commonly catalyze the oxidation of an organic substrate using a dioxygen molecule. The AlkB protein in plant RNA viruses is involved in the repair of methylation damage (van den Born et al. 2008). The host plant may use methylation as a defense mechanism to inactivate viral RNAs. Therefore, it has been suggested that AlkB homologues are involved in counteracting this defense mechanism (Bratlie & Drabløs 2005). However, selection for maintenance of the conserved *AlkB* domain could also involve environmental factors, such as the use of alkylating pesticides. *AlkB* domains are mainly found in plant viruses belonging to the *Flexiviridae* family, which are – like TEV – positive-sense single-stranded RNA viruses. Within the *Potyviridae* family, only one virus with an *AlkB* domain has been reported: *Blackberry virus Y* (BVY) (Susaimuthu et al. 2008). In BVY *AlkB* is located within the *P1* serine protease gene. Interestingly, BVY lacks the N-terminus of HC-Pro that is present in the potyvirus orthologs. This missing region is involved in silencing suppression and vector transmission in other potyviruses (Revers et al. 1999; Urcuqui-Inchima et al. 2001; Young et al. 2007; Stenger et al. 2006). These are important functions for a potyvirus, suggesting that AlkB has evolved to take over these functions in BVY. To test this idea in a model system, we have introduced the conserved AlkB domain within the P1 protein in TEV.

Whereas the newly introduced *AlkB* domain was rapidly purged from the TEV genome in our experimental setup, the functionally redundant *2b* gene was maintained for more than half a year of evolution. We then performed further experiments to show that *2b* is functional within the TEV genome, and even might be beneficial when the functional domain of HC-Pro related to RNA-silencing suppression is mutated.

## Materials and Methods

### Viral constructs, virus stocks and plant infection

The TEV genome used to generate the virus constructs was originally isolated from *Nicotiana tabacum* plants (Carrington et al. 1993). In this study two different variants of TEV containing exogenous sequences were used. One of these virus variants, hereafter named TEV-2b, contains the *2b* gene from CMV. The other virus variant, hereafter named TEV-AlkB, contains a conserved domain, characterized as a 2OG-Fe(II) oxygenase domain isolated from *Nicotiana benthamiana*.

TEV-2b and TEV-AlkB were generated from cDNA clones constructed using plasmid pMTEVa, which consists of a TEV infectious cDNA (accession: DQ986288, including two silent mutations, G273A and A1119G) flanked by SP6 phage RNA promoter derived from pTEV7DA (GenBank: DQ986288). pMTEVa contains a minimal transcription cassette to ensure a high plasmid stability (Bedoya & Daròs 2010). The clones were constructed using standard molecular biology techniques, including PCR amplification of cDNA with the high-fidelity Phusion DNA polymerase (Thermo Fisher Scientific), DNA ligation with T4 DNA ligase (Thermo Fisher Scientific) and transformation of *Escherichia coli* DH5α by electroporation. For generation of TEV-2b the *2b* gene was PCR amplified using phosphorylated primers forward 5’-ATGGAATTGAACGTAGGTGCA-3’ and reverse 5’-TTGAAAGTAGAGGTTCTCTGTAGTGAAAGCACCTTCCGCCC-3’ containing spacers and a proteolytic cleavage site between *2b* and *CP*. The *2b* gene was first ligated into the smaller pTV3 plasmid that was opened with primers: forward 5’-TCAGGTACAGTGGGTGCTGGTGTTGAC-3’ and reverse 5’-CTGAAAATAAAGATTCTCAGTCGTTG-3’. The restriction enzymes *Sal*I and *Bgl*II were used to transfer the digestion product containing the *2b* gene to the pMTEVa plasmid. For generation of TEV-AlkB, RNA was isolated from *N. benthamiana* and subsequently the conserved *AlkB* domain from the alkylated DNA repair protein AlkB homologue 1 was amplified by RT-PCR with primers forward 5’-GCCGCGCGCTCTGGAGCACCCTTGGCTTGCAGTTTG-3’ and reverse 5’-GCGCGTCTCTTGCCATGGGTGAATACTTGTCTAATGTTGATATTG-3’. After A-tailing with Taq DNA polymerase (Roche), the PCR product was cloned into pTZ57R/T (Thermo Fisher Scientific). Restriction sites for *Esp*3I and *Pau*I were then used for insertion of the cloned *AlkB* domain into pMTEVa. Sanger sequencing confirmed the sequences of the resulting plasmids.

The plasmids of TEV-2b and TEV-AlkB were linearized by digestion with *Bgl*II prior to *in vitro* RNA synthesis using the mMESSAGE mMACHINE® SP6 Transciption Kit (Ambion), as described in Carrasco et al. (2007). The third true leaf of 4-week-old *N. tabacum* L. cv Xanthi *NN* plants was mechanically inoculated with 5 μg of transcribed RNA. All symptomatic tissue was collected 7 days post-inoculation (dpi) and stored at −80 ºC as a viral stock.

### Experimental evolution

For the serial passage evolution experiments, 500 mg homogenized stock tissue was ground into fine powder using liquid nitrogen and a mortar, and resuspended in 500 μl phosphate buffer (50 mM KH_2_PO_4_, pH 7.0, 3% polyethylene glycol 6000). From this mixture, 20 μl were then mechanically inoculated on the third true leaf of 4-week old *N. tabacum* plants. At least five independent replicates were performed for each virus variant. At the end of the designated passage duration (3 or 9 weeks) all leaves above the inoculated leaf were collected and stored at −80 ºC. For subsequent passages the frozen tissue was homogenized and a sample was ground and resuspended with an equal amount of phosphate buffer (Zwart et al. 2014). Then, new *N. tabacum* plants were mechanically inoculated as described above. The plants were kept in a BSL-2 greenhouse at 24 ºC with 16 h light:8 h dark photoperiod.

### Reverse transcription polymerase chain reaction (RT-PCR)

To determine the stability of the exogenous sequences, RNA was extracted from 100 mg homogenized infected tissue using the InviTrap Spin Plant RNA Mini Kit (Stratec Molecular). Reverse transcription (RT) was performed using MMuLV reverse transcriptase (Thermo Fisher Scientific) and the reverse primer 5’-CGCACTACATAGGAGAATTAG-3’ located in the 3’UTR of the TEV genome. PCR was then performed with Taq DNA polymerase (Roche) and primers flanking the *2b* gene: forward 5’-TACGATATTCCAACGACTG-3’ and reverse 5’-GCAAACTGCTCATGTGTGG-3’, or the AlkB domain: forward 5’-GCAATCAAGCATTCTACTTC-3’ and reverse 5’-ATGGTATGAAGAATGCCTC-3’. PCR products were resolved by electrophoresis on 1% agarose gels.

### Fitness assays

The genome equivalents per 100 mg of tissue of the ancestral virus stocks and all evolved lineages were determined prior to subsequent fitness assays. The InviTrap Spin Plant RNA Mini Kit (Stratec Molecular) was used to isolate total RNA from 100 mg homogenized infected tissue. Real-time quantitative RT-PCR (RT-qPCR) was performed using the One Step SYBR PrimeScript RT-PCR Kit II (Takara), in accordance with manufacturer instructions, in a StepOnePlus Real-Time PCR System (Applied Biosystems). Specific primers for the CP gene were used: forward 5’-TTGGTCTTGATGGCAACGTG-3’ and reverse 5’-TGTGCCGTTCAGTGTCTTCCT-3’. The StepOne Software v.2.2.2 (Applied Biosystems) was used to analyze the data. The concentration of genome equivalents per 100 mg of tissue was then normalized to that of the sample with the lowest concentration, using phosphate buffer.

For the accumulation assays, 4-week-old *N. tabacum* plants were inoculated in the third true leaf with 50 μl of the normalized dilutions of ground tissue. For each ancestral and evolved lineage, at least three independent plant replicates were used. Leaf tissue was harvested 7 dpi. Total RNA was extracted from 100 mg of homogenized tissue. Virus accumulation was then determined by means of RT-qPCR for the *CP* gene of the ancestral and the evolved lineages. For each of the harvested plants, at least three technical replicates were used in the RT-qPCR.

To measure within-host competitive fitness, we used TEV carrying an enhanced green fluorescent protein (TEV-eGFP) (Bedoya & Daròs 2010) as a common reference competitor. TEV-eGFP has proven to be stable up to six weeks (using 1- and 3-week serial passages) in *N. tabacum* (Zwart et al. 2014), and is therefore not subjected to appreciable eGFP loss during our 1-week competition experiments. All ancestral and evolved viral lineages were again normalized to the sample with the lowest concentration, and 1:1 mixtures of viral genome equivalents were made with TEV-eGFP. The mixture was mechanically inoculated on 4-week-old *N. tabacum* plants, using three independent plant replicates per viral lineage. The plant leaves were collected at 7 dpi, and stored at −80 ºC. Total RNA was extracted from 100 mg homogenized tissue. RT-qPCR for the *CP* gene was used to determine total viral accumulation, and independent RT-qPCR reactions were also performed for the *eGFP* gene sequence using primers forward 5’- CGACAACCACTACCTGAGCA-3’ and reverse 5’-GAACTCCAGCAGGACCATGT-3’. The ratio of the evolved and ancestral lineages to TEV-eGFP is then, where 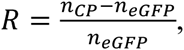 where *n_CP_* and 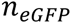 are the RT-qPCR measured copy numbers of *CP* and *eGFP*, respectively. Within-host competitive fitness can then be estimated as 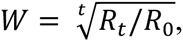, where *R*_0_ is the ratio at the start of the experiment and the ratio after *t* days of competition (Carrasco et al. 2007). Note that the method for determining only works well when the frequency of the common competitor is below ~0.75. This limitation was not problematic though, since in these experiments the fitness of the evolved virus populations remained the same or increased. The statistical analyses comparing the fitness between lineages were performed using R v.3.2.2 (R Core Team 2014) and IBM SPSS version 23.

### Generation and infectivity of the HC-Pro FINK and AS13 mutant genotypes

The plasmids of TEV and TEV-2b were used to generate two different mutant genotypes in each of the background sequences. One mutant genotype, hereafter named FINK, consists of the G548U nucleotide substitution within *HC-Pro*, which results in the R183I amino acid change within the highly conserved FRNK box. The other mutant genotype, hereafter named AS13, consists of the A895C and A900C nucleotide substitutions within *HC-Pro*, which results in the E299A and D300A amino acid changes. The TEV and TEV-2b FINK mutant genotypes were generated by site-directed mutagenesis performed with the QuickChange II XL Site-Directed Mutagenesis Kit (Agilent Technologies) on the original pMTEVa and pMTEVa-2b plasmids (described above). The primers used for mutagenesis are: forward 5’- GCGCATTGGCCACCTTGGTTCTTTCATAAATAAAATCTCATCGAAGGCCCATGTG-3’ and reverse 5’- CACATGGGCCTTCGATGAGATTTTATTTATGAAAGAACCAAGGTGGCCAATGCGC-3’. To obtain the TEV and TEV-2b AS13 mutant genotypes, the *HC-Pro* gene of TEV was replaced by that of the AS13 mutant. For this, an overlapping PCR product of ~3 kb was generated, composed of: a 5’ fragment amplified with primers forward 5’-GGTGGAGCTCGTATGGCTTGC-3’ and reverse 5’-GTCAATCTCAGTGAAAAATCCTTTAAGAAACC-3’; a central sequence amplified with primers forward 5’-GGGGCCGATCTCGAAGAGGCAAGCACAC-3’ and reverse 5’- TCCTGTCGTTCTAGACCCATACGAATCC-3’; and a 3’ fragment amplified with primers forward 5’-GATAATGTCAACTGACTTCCAGACGCTCAGGC-3’ and reverse 5’-AGAATGTCAACAACAAGGAAAGGACACTCG-3’. The PCR products were digested with *SnaB*I and *Aat*II enzymes and then ligated into the pMTEVa and pMTEVa-2b plasmids previously opened with the same enzymes. Sanger sequencing confirmed the sequences of the resulting plasmids.

To test the viability of TEV^FINK^, TEV-2b^FINK^, TEV^AS13^, and TEV-2b^AS13^ the third true leaf of 4-week-old *N. tabacum* plants was mechanically inoculated with equal amounts of transcribed RNA. Two different assays were done. In a first one, plants were left to grow 20 dpi to observe the development of symptoms. In a second assay, all tissue was collected 10 dpi and stored at −80 ºC. RNA was extracted from 100 mg homogenized tissue using the Agilent Plant RNA Isolation Mini Kit (Agilent Technologies). As many plants did not show symptoms, RT-qPCR for the *CP* gene was performed to confirm infection and to determine viral accumulation levels of the different mutant genotypes. For those plants that were infected with sufficient viral accumulation levels to perform RT-PCR, Sanger sequencing was performed to confirm the sequences of the FINK and AS13 genotypes. RT was performed using MMuLV reverse transcriptase (Thermo Fisher Scientific) and the reverse primer 5’- TTGCACCTTGTGTGACCAC-3’, located in the *P3* gene. PCR was then performed with Phusion DNA polymerase (Thermo Fisher Scientific) and primers flanking the region of interest: forward 5’-GGGGCCGATCTCGAAGAGGCAAGCACAC-3’ and reverse 5’-GCAAAGAAAATGTTCATGTAGCAATAACCTTCATTAGCTATATACATT-3’. Sanger sequencing was performed at GenoScreen (Lille, France: www.genoscreen.com; last accessed June 23, 2016) with an ABI3730XL DNA analyzer. Sequences were assembled using Geneious v.9.0.2 (www.geneious.com; last accessed June 23, 2016).

### Measurement of the relative expression of *NtAlkB*

The expression of the *AlkB* endogenous gene present in *N. tabacum*, hereafter referred to as *NtAlkB*, was measured by RT-qPCR. This measurement was done relative to two different reference genes: *L25*, encoding the ribosomal protein L25, and *EF1α*, encoding the translation elongation factor 1α (Schmidt & Delaney 2010). Total RNA was extracted from healthy plants and from plants infected with TEV and TEV-AlkB, respectively, at 7 dpi using three plant replicates each. To ensure that only the endogenous *NtAlkB* was amplified from TEV-AlkB- infected plants, primers outside the conserved domain were used. These primers were: forward 5’-TTCTGCTTGGAGAGTCATCCAG-3’ and reverse 5’-TCTCCTTTATCCATCGGCACTG-3’. The *L25* reference gene was amplified using forward primer 5’- CCCCTCACCACAGAGTCTGC-3’ and reverse primer 5’- AAGGGTGTTGTTGTCCTCAATCTT-3’. The *EF1α* gene was amplified with forward and reverse primers 5’-TGAGATGCACCACGAAGCTC-3’ and 5’- CCAACATTGTCACCAGGAAGTG-3’, respectively.

### Illumina sequencing, variants, and SNP calling

For Illumina next-generation sequencing (NGS) of the evolved and ancestral lineages, the viral genomes were amplified by RT-PCR using AccuScript Hi-Fi (Agilent Technologies) reverse transcriptase and Phusion DNA polymerase (Thermo Fisher Scientific), with six independent reactions that were pooled. Each virus was amplified using three primer sets generating three amplicons of similar size (set 1: 5’-GCAATCAAGCATTCTACTTCTATTGCAGC-3’ and 5’- TATGGAAGTCCTGTGGATTTTCCAGATCC-3’; set 2: 5’-TTGACGCTGAGCGGAGTGATGG-3’ and 5’-AATGCTTCCAGAATATGCC-3’; set 3: 5’- TCATTACAAACAAGCACTTG-3’ and 5’-CGCACTACATAGGAGAATTAG-3’). Equimolar mixtures of the three PCR products were made. Sequencing was performed at GenoScreen. Illumina HiSeq2500 2×100bp paired-end libraries with dual-index adaptors were prepared along with an internal PhiX control. Libraries were prepared using the Nextera XT DNA Library Preparation Kit (Illumina Inc.). Sequencing quality control was performed by GenoScreen, based on PhiX error rate and Q30 values.

Read artifact filtering and quality trimming (3’ minimum Q28 and minimum read length of 50 bp) was done using FASTX-Toolkit v.0.0.14 (http://hannonlab.cshl.edu/fastx_toolkit/index.html, last accessed April 10, 2016). De-replication of the reads and 5’ quality trimming requiring a minimum of Q28 was done using PRINSEQ-lite v.0.20.4 (Schmieder & Edwards 2011). Reads containing undefined nucleotides (N) were discarded. As an initial mapping step, the evolved sequences were mapped using Bowtie v.2.2.6 (Langmead & Salzberg 2012) against their corresponding ancestral sequence: TEV (GenBank accession number KX137149), TEV-2b ancestral (GenBank accession number KX832616), and TEV-AlkB ancestral (GenBank accession number KX832617). Subsequently, mutations were detected using SAMtools’ mpileup (Li et al. 2009) in the evolved lineages as compared to their ancestral lineage. At this point, we were only interested in mutations at a frequency > 10% and therefore, we present frequencies as reported by SAMtools, which has a low sensitivity for detecting low-frequency variants (Spencer et al. 2014).

After the initial pre-mapping step, error correction was done using Polisher v2.0.8 (available for academic use from the Joint Genome Institute) and consensus sequences were defined for every lineage. Subsequently, the cleaned reads were remapped using Bowtie v.2.2.6 against the corresponding consensus sequence for every lineage. For each new consensus, single nucleotide polymorphisms (SNPs) within each virus population were identified using SAMtools’ mpileup and VarScan v.2.3.9 (Koboldt et al. 2012). For SNP calling maximum coverage was set to 40000 and SNPs with a frequency < 1% were discarded.

## Results and Discussion

### Introducing functional redundancy in TEV genome: the *2b* VSR

To simulate HGT between two virus species, we have introduced the *2b* VSR from CMV into the TEV genome (Figure 1A), between the *NIb* replicase gene and the *CP* (Figure 1B). The C-terminus of 2b was modified to provide a proteolytic cleavage site for NIa-Pro similar to that between NIb and CP in wild-type TEV. The introduction of *2b* generates functional redundancy within TEV, as the multifunctional HC-Pro protein also acts as a silencing suppressor. We postulated that the functionally redundant TEV-2b virus could go down different trajectories, depending on (*i*) the fitness effect of *2b* upon insertion into the TEV genome, (*ii*) whether or not the gene can be accommodated in the genome, and (*iii*) whether *2b* could be maintained because functional redundancy increases the mutational robustness of TEV, buffering the effects of deleterious mutations hindering the silencing suppressor activity. If the insertion of *2b* is immediately beneficial, or if accommodation of the gene into the TEV genome occurs quickly, *2b* could be maintained over evolutionary time. In this scenario, it is possible that 2b becomes entirely responsible for silencing suppression, taking over from HC-Pro and allowing it to specialize on one of its other functions.

**Fig 1.**
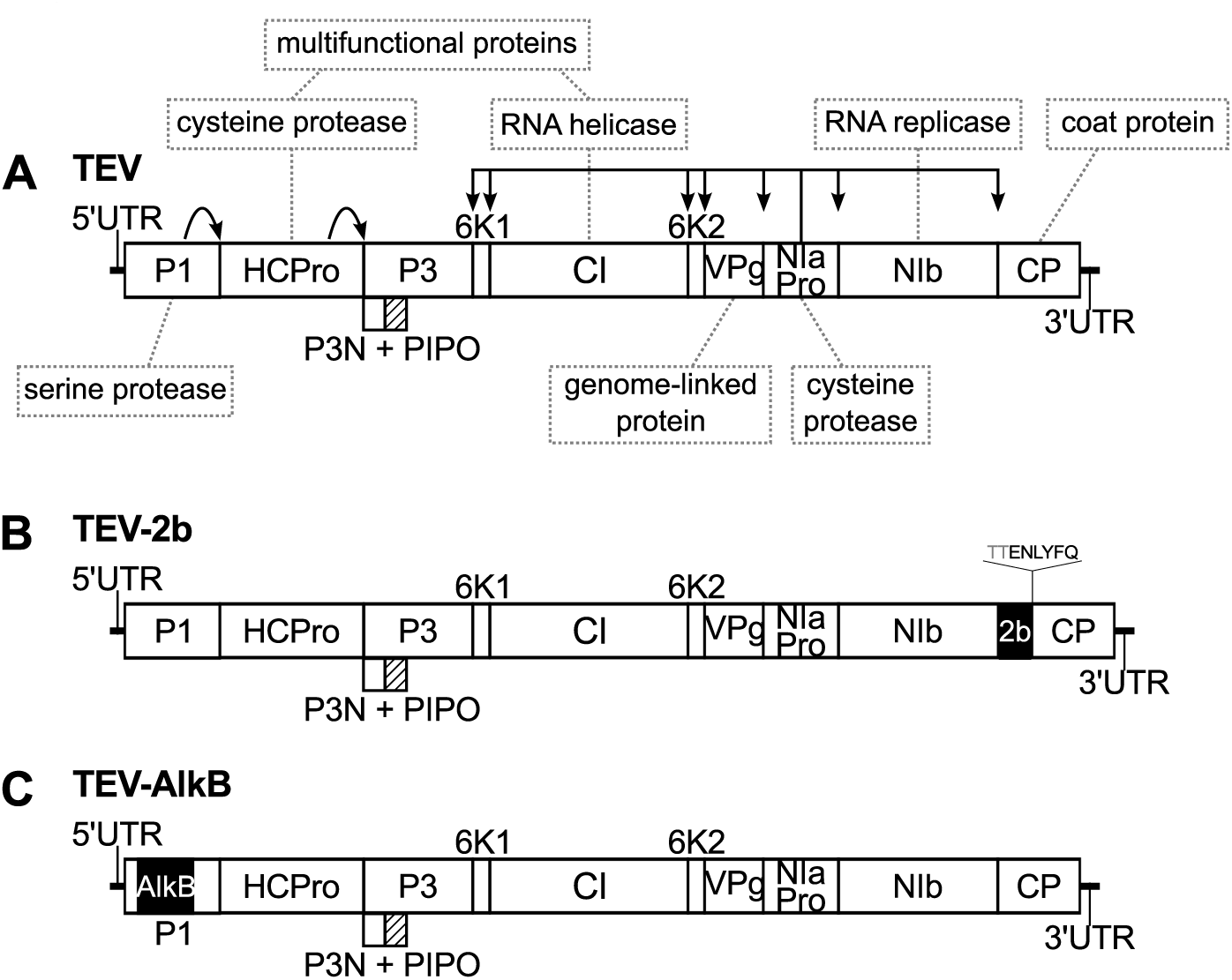
Schematic representation of the *Tobacco etch virus* genotypes with exogenous gene insertions. (A) The wild-type TEV codes for 11 mature peptides, including P3N-PIPO embedded within the P3 protein at a +2 frameshift. Relevant names of the protein products are indicated in the grey dotted boxes. (B) The *2b* gene from *Cucumber mosaic virus* was introduced between *NIb* and *CP*. The C-terminus of *2b* was modified to provide a similar proteolytic cleavage site for NIa-Pro as the one that originally existed between *NIb* and *CP*. (C) A conserved *AlkB* domain, similar to the one found in the host plant, was introduced within the *P1* gene.

When we evolved TEV-2b virus in *N. tabacum* plants, using 3-week and 9-week passages, we never detected any genomic deletions by RT-PCR. These results contrasts with all previous long-duration passage results for TEV, where deletions removing the added genetic material were quickly observed, regardless of it being a redundant gene copy (Willemsen, et al. 2016a; Willemsen, et al. 2016b) or encoding for additional marker proteins (Dolja et al. 1993; Zwart et al. 2014). Therefore, we speculated that the TEV-2b virus might have a fitness advantage over the wild-type TEV. To test this idea, we measured within-host competitive fitness and viral accumulation of the ancestral and evolved lineages. Additionally, we measured plant height as a proxy for virulence; healthy plants grow taller compared to plants infected with wild-type TEV. We found that the ancestral TEV and TEV-2b have a very similar within-host competitive fitness (Figure 2A; *t*-test comparing ancestral viruses: *t*_4_ = 2.008, *P* = 0.115), viral accumulation (Figure 2B; *t*-test comparing ancestral viruses: *t*_4_ = 1.389, *P* = 0.237), and plants grow to similar heights when infected with either of these viruses (supplementary Figure S1, Supplementary Material online; *t*-test comparing plants infected with ancestral viruses: *t*_4_ = 0.447, *P* = 0.678). Comparing the evolved lineages, the within-host competitive fitness of the TEV-2b lineages is similar to the lineages of TEV (Figure 2A; Mann-Whitney *U* = 4, *P* = 0.095). In addition, no significant difference in accumulation (Figure 2B; Mann-Whitney *U* = 18, *P* = 0.310) or plant height (supplementary Figure S1, Supplementary Material online; Mann-Whitney *U* = 8.5, *P* = 0.462) were found between the evolved lineages. All plants infected with TEV or TEV-2b were significantly shorter in height compared to the healthy control plants (supplementary Figure S1, Supplementary Material online; *t*-test with Holm-Bonferroni correction for multiple tests). These results show that the inserted *2b* gene has no effect on the fitness of TEV.

**Fig 2.**
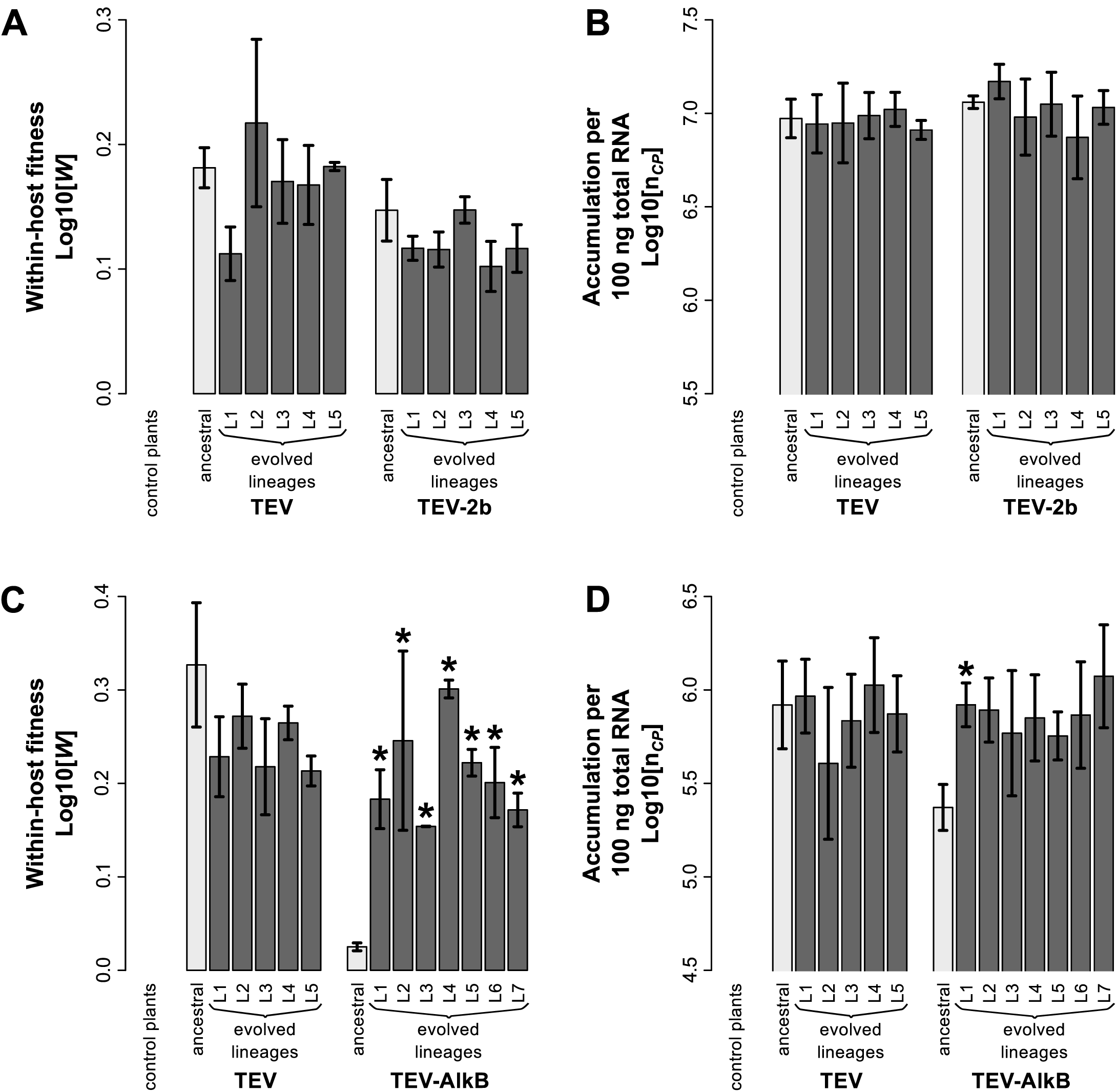
Fitness assays of the ancestral and evolved lineages. (A) Within-host competitive fitness (*W*), as determined by competition experiments and RT-qPCR, of TEV and TEV-2b with respect to a common competitor; TEV-eGFP. (B) Virus accumulation (*n_CP_*), as measured by RT-qPCR, of TEV and TEV-2b. (C) Within-host competitive fitness of TEV and TEV-AlkB. (D) Virus accumulation of TEV and TEV-AlkB. The ancestral lineages are indicated with light-gray bars and the evolved lineages with dark-gray bars. Both TEV and TEV-2b were evolved for a total of 27 weeks (three 9-week passages), using 5 replicate lineages for TEV and TEV-2b (L1- L5) and 7 replicate lineages (L1-L7) for TEV-AlkB. Evolved lineages that tested significantly different compared to their ancestral virus are indicated with an asterisk (*t*-test with Holm-Bonferroni correction for multiple tests).

All evolved and ancestral lineages were fully sequenced by Illumina technology. The sequences of the ancestral lineages were used as an initial reference for mapping of the evolved lineages. When looking at the genome sequences of the evolved TEV-2b lineages, we find no evidence of convergent adaptive evolution other than for mutations that are also present in the evolved wild-type TEV lineages (Figure 3A-B). Nonsynonymous mutation *NIb*/U7262C (amino acid replacement I94T) that appears in 2/10 of the evolved wild-type lineages has not been observed in any of the evolved TEV-2b lineages, suggesting a possible genotype-dependent fitness effect.

**Fig 3.**
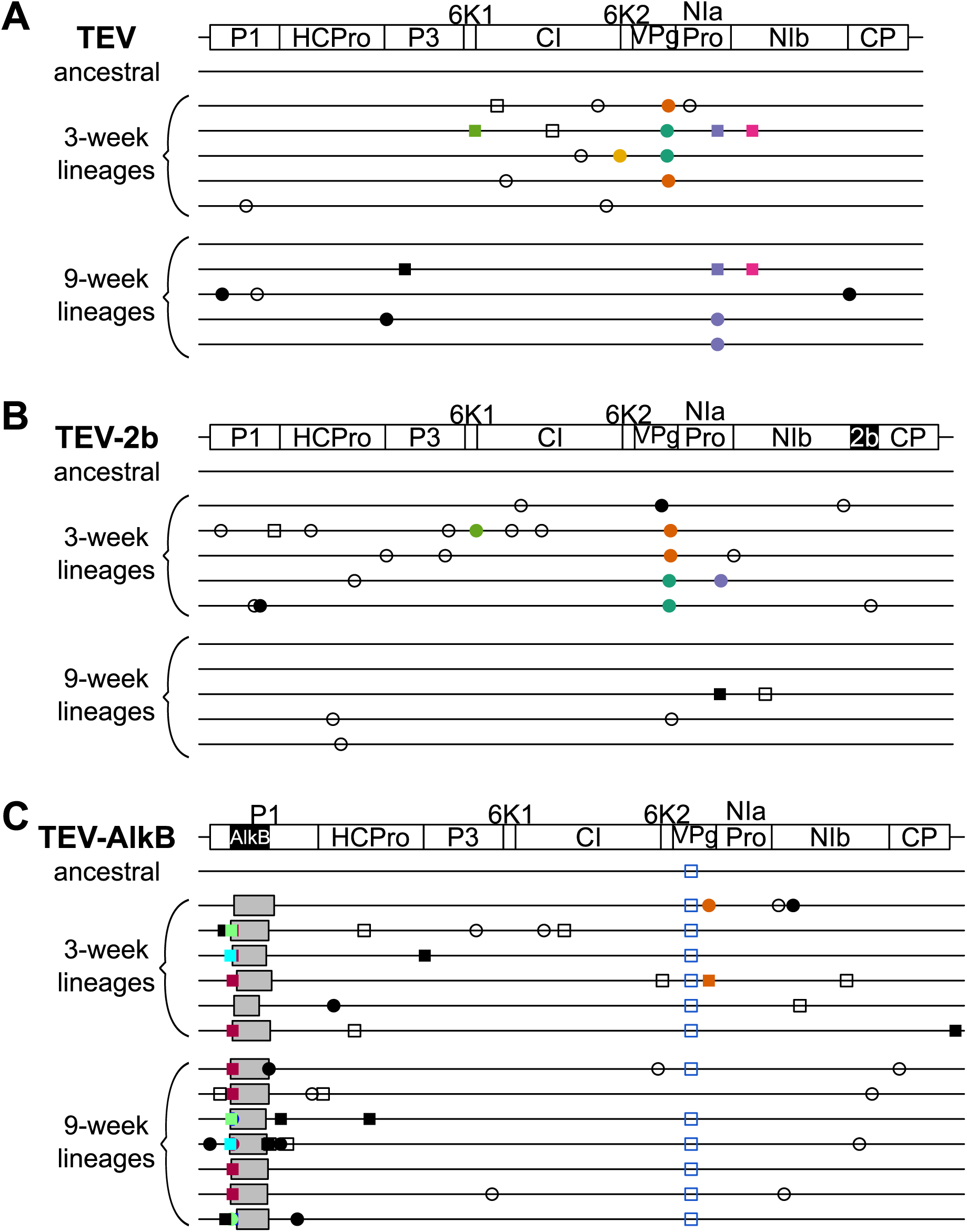
Genomes of the ancestral and evolved lineages. Mutations were detected using NGS data of the evolved virus lineages as compared to their ancestral lineages. The wild-type TEV is given for comparative purposes. The names on the left identify the virus genotypes and the lineages with their corresponding passage length. The square symbols represent mutations that are fixed (> 50%) and the circle symbols represent mutations that are not fixed (< 50%). Filled symbols represent nonsynonymous substitutions and open symbols represent synonymous substitutions. Black symbols represent substitutions occurring only in one lineage, whereas colored symbols represent substitutions that appear in two or more lineages, or in a lineage from another virus genotype. Note that the mutations are present at different frequencies as reported by SAMtools. Grey boxes indicate genomic deletions in the majority variant.

After remapping the cleaned reads against a newly defined consensus sequence for each lineage, we looked at the variation within each lineage. SNPs were detected from a frequency as low as 1%. In the evolved TEV-2b lineages a total of 507 (390 unique) SNPs were detected, of which 403 were found in the 3-week lineages and 104 in the 9-week lineages, with a median of 30.5 (range 17-149) per lineage. In the evolved TEV lineages a total of 379 (326 unique) SNPs were detected, of which 188 were found in the 3-week lineages and 191 in the 9-week lineages, with a median of 35 (range 17-63) per lineage. The distribution of SNPs per gene of the TEV-2b 3-week lineages is different compared to that of the wild-type 3-week lineages (Wilcoxon signed rank test; *V* = 0, *P* = 0.004), where in TEV-2b more SNPs accumulate in all genes (except for *VPg*, where an equal number of SNPs was found). By contrast, the 9-week lineages of TEV-2b accumulated less SNPs in most genes (all except *6K1* and *NIa-Pro*) than the wild-type lineages (Wilcoxon signed rank test; *V* = 61, *P* = 0.014). Interestingly, there is an inverse shift in the amount of synonymous versus nonsynonymous SNPs when comparing the TEV-2b 3-week (40% synonymous and 60% nonsynonymous) and 9-week lineages (67% synonymous and 33%: nonsynonymous) (Fisher’s exact test: *P* < 0.001), while this shift is not observed in the TEV 3- week (66% synonymous and 34% nonsynonymous) and 9-week (60% synonymous and 40% nonsynonymous) lineages (Fisher’s exact test: *P* = 0.464). These differences between the 3-week and 9-week lineages may result from the selective accommodation of the *2b* gene, as we previously observed that selection acts more strongly in the longer-duration passages (Zwart et al. 2014). For an overview of the SNP frequency per nucleotide position within the evolved lineages, see supplementary Figure S2, Supplementary Material online. For more details on the frequency of the SNPs within every lineage, see supplementary Tables S1 and S2, Supplementary Material online.

The shift of TEV-2b to a similar ratio of synonymous versus nonsynonymous SNPs as the wild-type might indicate that *2b* is successfully being accommodated in the viral genome. However, an alternative explanation is that the fitness cost of *2b*, which is small in size, is so low that there is little selection for viruses with genomic deletions, and the insert would only be eventually removed by genetic drift. As shown previously, the stability of gene insertions in the TEV genome depends, among other factors, on the position of the insertion (Willemsen, et al. 2016b). However, the introduction of two non-functional genes, the colored markers *Ros1* and *eGFP*, at the same genomic position, did lead to instability and the deletion of these markers over evolutionary time (Majer et al. 2013).

Neither the genome sequences of the evolved populations, nor the fitness and virulence assays, provide evidence that *2b* is actually adaptively accommodated into TEV or performing a function. To directly test whether *2b* is functional and functionally duplicates the silencing suppression activity of HC-Pro when inserted in the TEV genome, we generated mutant genotypes with mutations affecting functional domains of HC-Pro related to RNA silencing suppression. Two different mutant genotypes were considered. One is the so-called FINK mutant, which has an amino acid change (R183I) in the FRNK box in the central region of HC-Pro. The FRNK box is a highly conserved motif within all potyviruses which is required for sRNA binding and silencing suppression, and this activity is correlated with symptom severity (Shiboleth et al. 2007; Wu et al. 2010). The R183I mutation affects viral pathogenicity and has led to a significant reduction in symptom severity and viral titers in other potyviruses (Gal-On 2000; Lin et al. 2007; Shiboleth et al. 2007; Kung et al. 2014). The other mutant genotype, AS13, has the E299A and D300A amino acid substitutions (Kasschau et al. 1997). This mutant has reduced suppressor activity, low accumulation levels, and either does not induce symptoms at all or, in a few cases, mild mottle in *N. benthamiana* plants (Torres-Barceló et al. 2008, 2010).

After generating the FINK and AS13 variants in the TEV and TEV-2b background sequences, we performed infectivity assays in *N. tabacum* plants and scored symptoms at different time points (Figure 4). At 7 days post-inoculation (dpi) all plants infected with TEV, TEV-2b, TEV-2b^FINK^ and TEV-2b^AS13^ viruses display symptoms. The TEV-2b^FINK^ and TEV- 2b^AS13^ induce wild-type virus like symptoms in all of the infected plants. Most of the plants infected with TEV^FINK^ and TEV^AS13^ looked similar to healthy plants, although some plants (TEV^FINK^: 4/15; TEV^AS13^: 2/15) did show mild vein clearing in the upper leaves. When comparing the mean time to infection by means of a Kaplan-Meier log-rank test, the symptoms of TEV-2b^AS13^ appear significantly later compared to the wild-type TEV and TEV-2b viruses (Table 1). The symptoms of TEV-2b^FINK^ only appear significantly later compared to the wild-type TEV, but not compared to TEV-2b (Table 1). Nevertheless, the difference in the average time to infection between the wild-type TEV and TEV-2b is not significant (Table 1).

**Fig 4.**
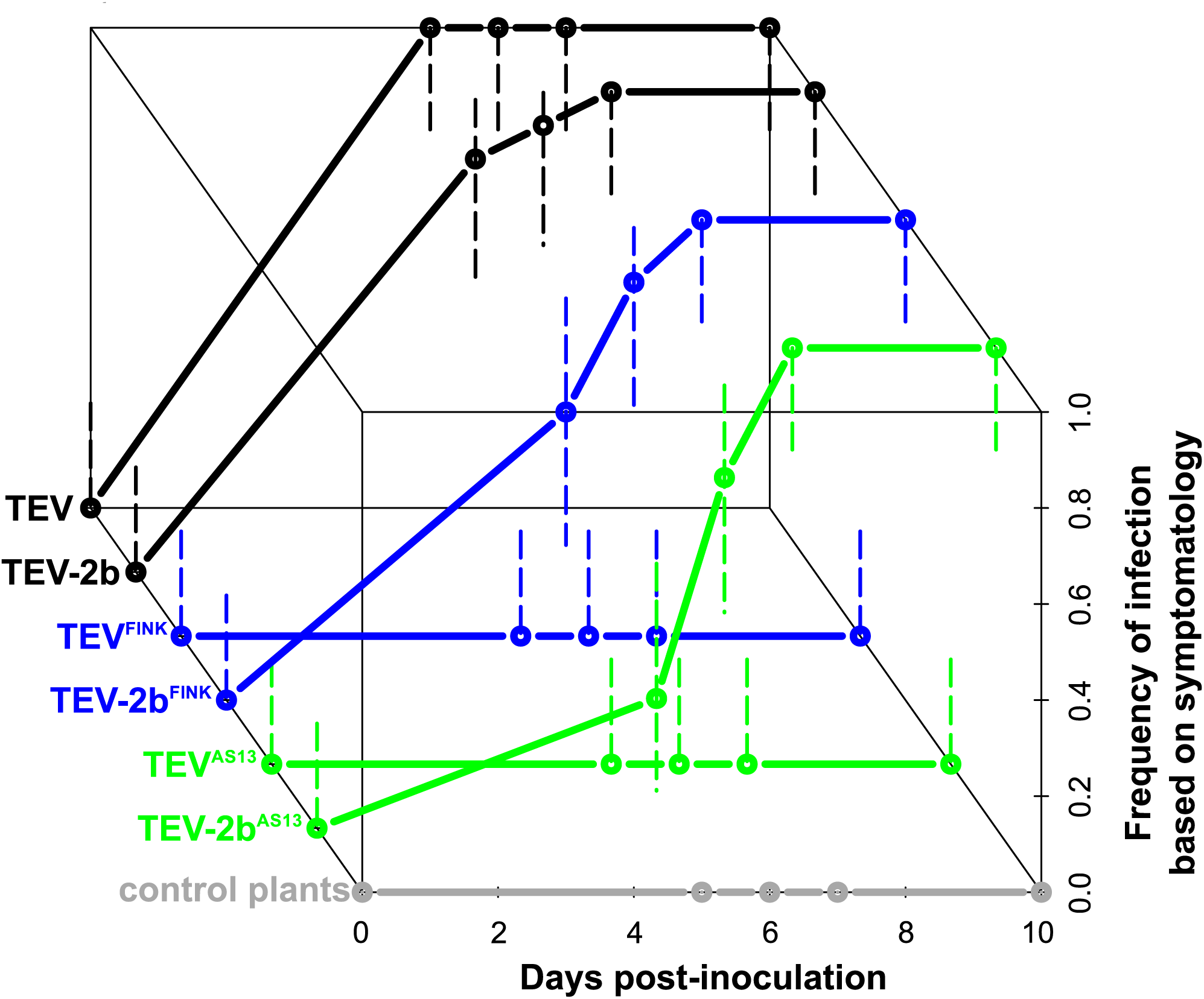
Frequency of symptomatic plants infected with the FINK and AS13 virus variants. The *z*-axis represents the cumulative frequency of infection of the TEV^FINK^, TEV-2b^FINK^, TEV^AS13^, and TEV-2b^AS13^ variants in *N. tabacum* plants, as compared to the wild-type TEV and TEV-2b viruses. The different viral genotypes are indicated on the ordinate. Error bars represent the 95% confidence intervals of the estimated frequencies.

**Table 1.** Kaplan-Meier Log-Rank Test of the FINK and AS13 infectivity data.

| Genotype | TEV |  | TEV-2b |  | TEV-2b<sup>FINK</sup> |  | TEV-2b<sup>AS13</sup> |  |
| --- | --- | --- | --- | --- | --- | --- | --- | --- |
| | $\chi^2$ | *P* | $\chi^2$ | *P* | $\chi^2$ | *P* | $\chi^2$ | *P* |
| TEV |  |  |  |  |  |  |  |  |
| TEV-2b | 2.071 | 0.150 |  |  |  |  |  |  |
| TEV-2b<sup>FINK</sup> | 7.250 | 0.007 | 1.596 | 0.206 |  |  |  |  |
| TEV-2b<sup>AS13</sup> | 16.789 | < 0.001 | 7.792 | 0.005 | 2.407 | 0.121 |  |  |

Even though TEV^FINK^ and TEV^AS13^ do not induce symptoms in *N. tabacum*, the possibility exists that plants are infected and that these viruses accumulate at low levels. Therefore, RT-PCR and RT-qPCR were used to test if asymptomatic plants inoculated with these two viruses were infected, to confirm the identity of the virus, and to measure accumulation. The HC-Pro region encompassing the FINK and AS13 mutations was amplified by RT-PCR and the mutant genotype of the positive samples was confirmed by Sanger sequencing. The presence or absence of the *2b* gene was also confirmed by amplifying the region encompassing *2b*. We found that 14/15 plants inoculated with TEV^FINK^ were infected with the virus, while only 4/15 plants inoculated with TEV^AS13^ were infected. The viral copy numbers in each plant were measured by RT-qPCR (Figure 5), and indeed, we found that TEV^FINK^ and TEV^AS13^ accumulate at very low levels. When comparing the RT-qPCR measured viral copy numbers of TEV^FINK^ and TEV- 2b^FINK^ as well as of TEV^AS13^ and TEV-2b^AS13^, we found a significant effect of the *HC-Pro* allele present and the plant replicate within each virus genotype (*i.e*., presence or absence of *2b*) (Table 2), indicating that the TEV-2b^FINK^ and TEV-2b^AS13^ accumulate at higher levels compared to the TEV^FINK^ and TEV^AS13^ viruses. By contrast, no significant difference was found when comparing the wild-type TEV and TEV-2b (Table 2).

**Fig 5.**
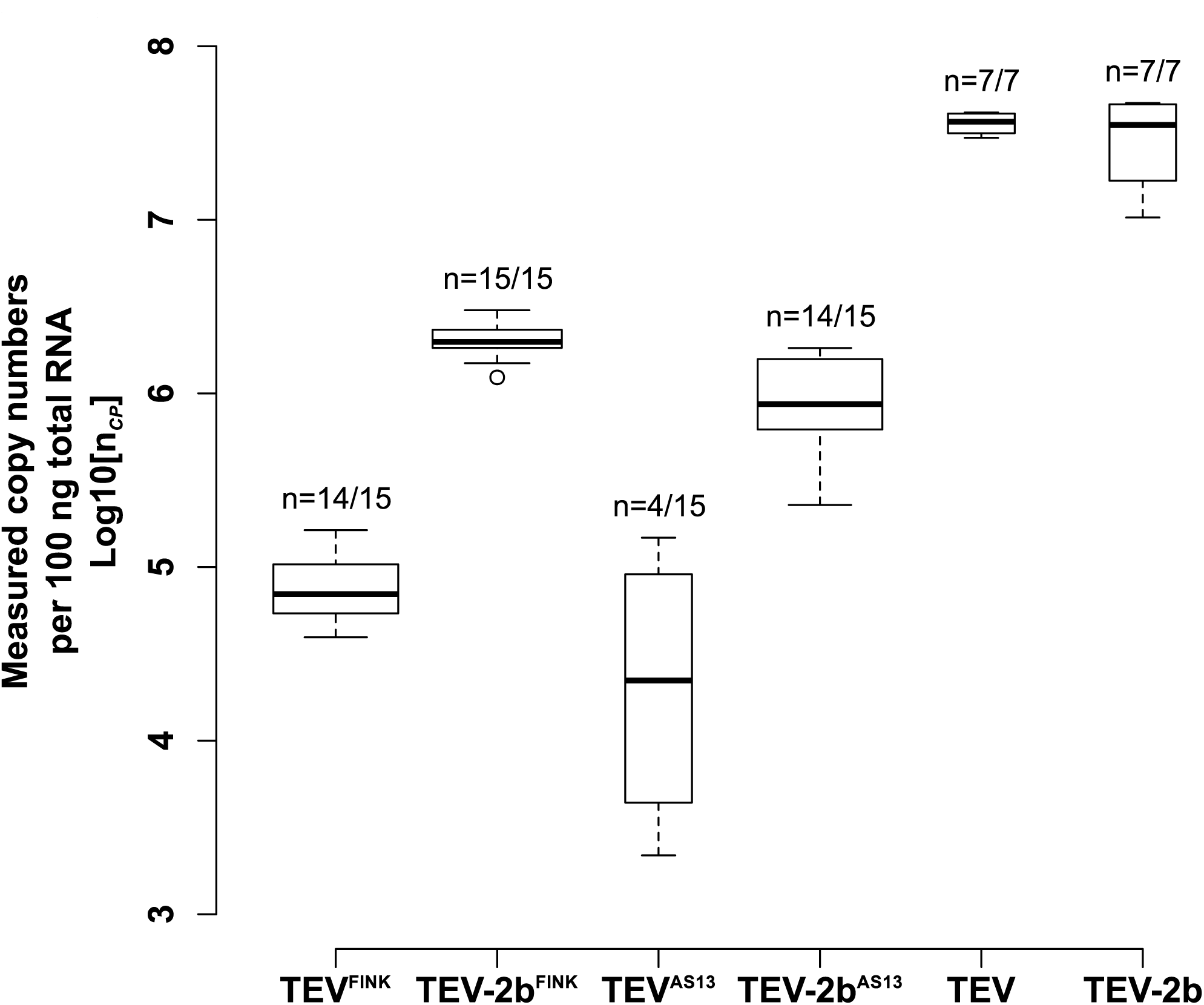
Measured level of infection of the FINK and AS13 variants. RT-qPCR measured virus accumulation (*n_CP_*) of the TEV^FINK^, TEV-2b^FINK^, TEV^AS13^, and TEV-2b^AS13^ variants. The wild-type TEV and TEV-2b viruses are given for comparative purposes. The numbers above the boxes indicate the number of plants infected over the total number of plants tested. Error bars represent the standard deviation of the mean, whilst circles represent outliers.

**Table 2.** Nested ANOVAs on RT-qPCR measured viral copy numbers.

| **Comparison** | **Source of Variation** | **SS** | **df** | **MS** | ***F*** | ***P*** |
| --- | --- | --- | --- | --- | --- | --- |
| TEV<sup>FINK</sup> vs TEV-2b<sup>FINK</sup> | Genotype | 44.634 | 1 | 44.634 | 735.467 | < 0.001 |
|  | Plant within Genotype | 1.639 | 27 | 0.061 | 145.065 | < 0.001 |
|  | Error | 0.024 | 58 | 0.000 |  |  |
| TEV<sup>AS13</sup> vs TEV-2b<sup>AS13</sup> | Genotype | 24.327 | 1 | 24.327 | 41.595 | < 0.001 |
|  | Plant within Genotype | 9.358 | 16 | 0.585 | 1743.434 | < 0.001 |
|  | Error | 0.012 | 36 | 0.000 |  |  |
| TEV vs TEV-2b | Genotype | 0.334 | 1 | 0.334 | 2.665 | 0.129 |
|  | Plant within Genotype | 1.503 | 12 | 0.125 | 302.633 | < 0.001 |
|  | Error | 0.012 | 28 | 0.000 |  |  |

The lack of symptoms observed for TEV^FINK^ is consistent with the drastic reduction of symptom severity observed for this mutation in *Zucchini yellow mosaic virus* (ZYMV) in various cucurbit species (Gal-On 2000). However, in contrast to our results the FINK mutation in ZYMV did not noticeably affect virus accumulation or infectivity (Gal-On 2000; Shiboleth et al. 2007). In addition, the HC-Pro suppressor activity was not diminished for the ZYMV^FINK^ (Shiboleth et al. 2007). The FINK mutation appears to have different effects on the HC-Pro suppressor activity as this mutation in TEV, *Turnip mosaic virus* (TuMV), and *Potato virus Y* (PVY) did inactivate the HC-Pro suppressor activity (Shiboleth et al. 2007). For the TEV^AS13^ mutant genotype similar results were obtained in *N. benthamiana* plants, where no symptoms and reduced accumulation levels were observed (Torres-Barceló et al. 2008, 2010). Interestingly, a *Wheat streak mosaic virus* (WSMV) genome lacking the whole *HC-Pro* gene was viable in wheat, oat and corn, producing typical symptoms on all three hosts, however with lower virus titers (Stenger et al. 2005). These results indicate that single point mutations can have a similar effect as deleting the whole *HC-Pro*.

These data show that small functional sequences that duplicate a viral function can be maintained in a viral RNA genome. Regardless of whether such a sequence is maintained because it has only a minor fitness cost, or because of second-order evolutionary benefits, functional redundancy therefore could be conducive to evolutionary innovation by ameliorating constraints to virus adaptation that arise due to the rugged topology of the fitness landscape. For example, either the original virus gene or the added gene traverses a fitness valley that had restricted evolution to a local adaptive peak, given that one of the two copies can accumulate deleterious mutations without immediately lowering viral fitness. Gene duplication is often suggested as a mechanism for overcoming such constraints, but for plant viruses gene duplications rarely have been observed. Moreover, duplicated genes – including *HC-Pro –* are quickly lost (Willemsen, et al. 2016b), suggesting that HGT of smaller sequences providing functional redundancy might be an interesting alternative mechanism.

### Introducing a novel function into TEV genome: the *AlkB* domain

Based on the relative position of the *AlkB* domain within the *P1* gene in BVY, we have introduced the a 2OG-Fe(II) oxygenase *AlkB* domain from *N. benthamiana* into the TEV genome (Figure 1C). The *N. benthamiana* and *N. tabacum AlkB* domains share 97% nucleotide identity. In an attempt to preserve predicted protein secondary structures, the conserved *AlkB* domain was introduced within the *P1* gene without any proteolytic cleavage sites. The TEV-AlkB virus was viable in *N. tabacum* plants, although the symptoms induced were very mild. The expression of *NtAlkB* was higher in uninfected plants, compared to plants infected with TEV (Figure 6; *t*-test reference gene *L25*: *t*_4_ = 9.520, *P* < 0.001; reference gene *EF-1α*: *t*_4_ = 27.592, *P* < 0.001) or TEV- AlkB (Figure 6; *t*-test reference gene *L25*: *t*_4_ = 4.767, *P* = 0.009; reference gene *EF-1α*: *t*_4_ = 13.863, *P* < 0.001). The plants infected with TEV-AlkB, however, expressed *NtAlkB* at a significantly higher level than plants infected with TEV (*t*-test reference gene *L25*: *t*_4_ = 3.305, *P* = 0.030; reference gene *EF-1α*: *t*_4_ = 8.775, *P* < 0.001).

**Fig 6.**
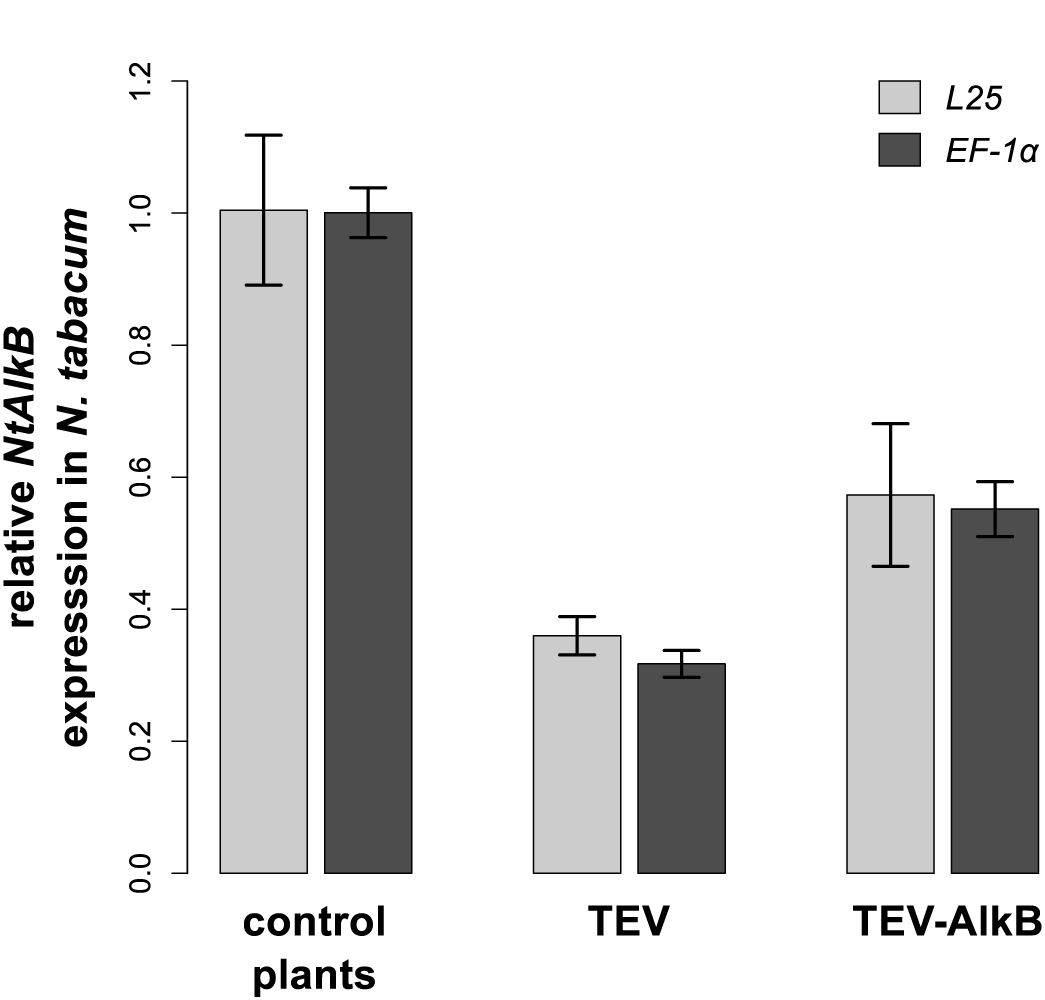
Relative *NtAlkB* expression of plants infected with TEV-AlkB. The expression of *NtAlkB* in *N. tabacum* plants was measured by RT-qPCR, relative to the two stable reference genes *L25* (ribosomal protein) and *EF-1α* (elongation factor 1α). Relative expression levels are given in healthy plants, plants infected with the wild-type TEV, and plants infected with TEV- AlkB. Light-gray bars indicate that *L25* was used as a reference, and dark gray bars indicate that *EF-1α* was used.

We then serially passaged the TEV-AlkB virus for 27 weeks, with each passage having a duration of either 3 or 9 weeks. We observed a variety of deletions already in the first passage, when RT-PCR amplifying the region encompassing the *AlkB* domain (Figure 7). A variety of small deletion variants were observed in the first 3-week passage (Figure 7A), while after a single 9-week passage there were apparently larger deletions (Figure 7B). At the end of the evolution experiment, all TEV-AlkB lineages contained large deletions within the *AlkB* domain, making it highly unlikely for this domain to remain functional.

**Fig 7.**
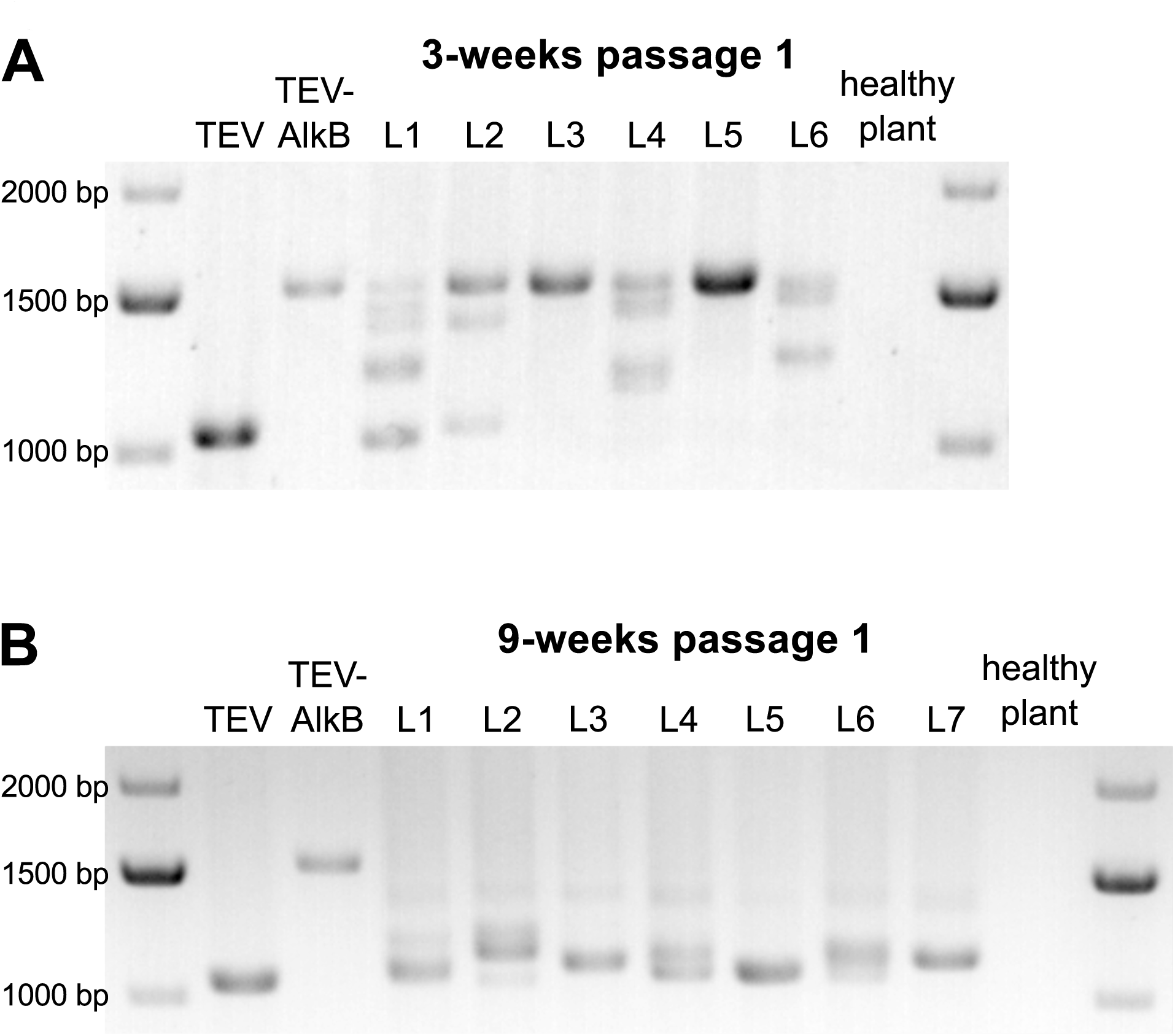
Detection of deletions after the first passage of TEV-AlkB. Agarose gels with RT-PCR products of the region encompassing the *AlkB* domain. Deletions were detected in lineages (L1-L6) after the first 3-week passage (A) and in lineages (L1-L7) of the first 9-week passage (B). In both gels the product sizes of the TEV and TEV-AlkB are shown as a reference, followed by the independent lineages, followed by a healthy control plant.

Next, we measured within-host competitive fitness in direct competition experiments with a fluorescently labeled TEV-eGFP, and we also measured viral accumulation. The introduction of the conserved *AlkB* domain within the TEV genome resulted in a significant decrease in within-host competitive fitness (Figure 2C; *t*-test comparing ancestral viruses: *t*_4_ = 7.846, *P* = 0.001) and viral accumulation (Figure 2D; *t*-test comparing ancestral viruses: *t*_4_ = 3.587, *P* = 0.023), compared to the wild-type TEV. After 27 weeks of evolution and the occurrence of partial or complete deletions of the *AlkB* domain, both within-host competitive fitness (Figure 2C; comparing evolved lineages: Mann-Whitney *U* = 10, *P* = 0.268) and viral accumulation (Figure 2D; comparing evolved lineages: Mann-Whitney *U* = 18, *P* = 1) were restored to levels similar to the evolved lineages of the wild-type virus.

All evolved and ancestral lineages were fully sequenced by Illumina technology. After an initial mapping step using the ancestral sequence as a reference, the positions of the majority deletions were defined (Figure 3C, Materials and Methods). Out of 13 evolved lineages, 8 lineages contain deletions within the inserted *AlkB* domain, of which 2 lineages contain an exact deletion of *AlkB*. For the remaining 5 lineages, deletions occurred either upstream (2/13 lineages) or downstream (3/13 lineages) the *AlkB* sequence, thus encompassing part of the *P1* serine protease gene. The P1 proteinase activity is not essential for viral infectivity (Verchot & Carrington 1995b), however the cleavage that separates P1 and HC-Pro is required (Verchot & Carrington 1995a). Based on TEV-AlkB infectivity and the fitness data of the evolved lineages, it does not seem that the proteolytic activity of P1 is affected by the deletions observed in this gene.

When mutations were detected in the sequenced lineages, evidence for convergent evolution was found (Figure 3C). A nonsynonymous mutation was found within the *AlkB* domain (U439C, amino acid replacement W10R) in 9/13 evolved lineages. However, in 8/9 lineages this mutation is present in the sequences of the minority deletion variants, in other words, this mutation falls within the deleted region of the majority deletion variant (Figure 3C) and is supported by a low read coverage. Another mutation was found in 12/13 evolved lineages in *VPg* (A6429C). This is a synonymous mutation and is fixed in all 12 lineages, however, this mutation was also found to be fixed within the ancestral population. For more details on the SNPs within every lineage with a frequency > 1%, see supplementary Table S3, Supplementary Material online.

The reduction of *NtAlkB* expression in the host plant when infected with TEV, suggests that TEV might have a mechanism to suppress the expression of this gene in the host plant. AlkB activity could be beneficial for the host plant as a mechanism of immunity against pathogens like TEV, as it is involved in methylation. The introduction of the conserved *AlkB* domain within the TEV genome does not result in complete silencing of the plant *NtAlkB* gene. On the contrary, TEV-AlkB does not suppress the expression of *NtAlkB* to similar low levels as the wild-type virus does. The higher *NtAlkB* expression in the host plant when infected with TEV-AlkB is likely to be an effect of slower replication of this virus. This is corroborated by the reduced symptomatology, within-host competitive fitness, and viral accumulation of TEV-AlkB. The pseudogenization of the *AlkB* domain confirms that – for this experimental setup – this gene is not beneficial for TEV, or that any benefits of *AlkB* are outweighed by the costs.

### Concluding remarks

In this study, we have simulated two independent events of HGT: from (*i*) an unrelated virus and from (*ii*) the host species into the genome of TEV. Both inserted genes could provide additional enzymatic activities to the resulting recombinant viruses. Moreover, in the first case, functional redundancy with one of the TEV’s own proteins was introduced. We explored the evolutionary fate of the resulting recombinant TEV genomes.

Overall, the results from the first HGT experiment strongly suggest that the *2b* gene from CMV can compensate for what would otherwise be deleterious mutations in HC-Pro. Therefore, it seems reasonable to conclude that 2b exhibits VSR activity as a transgene in TEV, to the extent that it can compensate for the hypo-suppression of the AS13 and FINK mutants. Even though symptoms develop more slowly, infection is similar to both the TEV-2b^FINK^ and TEV-2b^AS13^ genotypes 10 dpi. There are therefore two possible evolutionary mechanisms to explain these results. First, although the 2b protein has been shown to be functional in TEV-2b, it could still not be beneficial and at the same time have a small fitness cost. In this case, the gene would be maintained during the evolution experiment due to (*i*) a limited mutational supply of large genomic deletions that conserve polyprotein processing, (*ii*) weak selection for these mutants and (*iii*) periodic bottlenecks during horizontal transmissions that reduce the efficiency of selection of slightly beneficial deletions. Second, our results with the HC-Pro mutants show that the functional redundancy provided by 2b may, in principle, increase the mutational robustness of the VSR activity. Therefore, *2b* could be maintained because it contributes to buffering the effects of deleterious mutations in the silencing suppressor region of HC-Pro.

In contrast to the observed evolutionary stability of the *2b* gene, the simulated HGT of the *AlkB* domain from *Nicotiana* into the TEV genome turns out to be highly unstable. The *AlkB* domain sequence inserted has very low similarity to the *AlkB* domain found in BVY, the only other potyvirus that carries this domain. Furthermore, the *AlkB* domain in BVY has orthologous domains in other plant viruses (*e.g*., *Flexiviridae*), and the similarity between these is higher than the similarity of BVY P1 orthologs in other potyviruses (Susaimuthu et al. 2008). Therefore, it is more likely that BVY has obtained this domain through a HGT event from another virus rather than from its host. A similar experimental setup but with the *AlkB* domain from BVY inserted into TEV might potentially lead to different results, resulting in the maintenance and accommodation of this gene.

In conclusion, HGT events may contribute to increase the genome complexity of RNA viruses, but the evolutionary outcome of such events would hardly be predictable. Adding new functions does not necessarily result in retaining the novel gene.

## Acknowledgements

We thank Francisca de la Iglesia and Paula Agudo for excellent technical assistance.

## Supplementary Files

### Supplementary file 1

This file contains: supplementary Figure S1 with the plant height as a proxy for virus induced virulence of plants infected with TEV and TEV-2b; and supplementary Figure S2 with the SNP frequency per nucleotide position within the evolved TEV and TEV-2b viral lineages.

### Supplementary file 2

This file contains supplementary Tables S1, S2 and S3 with the within population sequence variation of the evolved and ancestral TEV, TEV-2b and TEV-AlkB lineages, respectively.

## Data Deposition

The sequences of the ancestral viral stocks were submitted to GenBank with accessions TEV: KX137149; TEV-2b: KX832616; TEV-AlkB: KX832617). The raw read data from Illumina sequencing is available at SRA with accession: SRP090412. The fitness data has been deposited on LabArchives with doi: 10.6070/H4DF6P97.

## Funding

This work was supported by the John Templeton Foundation (grant 22371), the European Commission 7^th^ Framework Program EvoEvo Project (grant ICT-610427), and the Spanish Ministerio de Economía y Competitividad (grants BFU2012-30805 and BFU2015-65037-P) to S.F.E. The opinions expressed in this publication are those of the authors and do not necessarily reflect the views of the John Templeton Foundation. The funders had no role in study design, data collection and analysis, decision to publish, or preparation of the manuscript.

## References

Aravind L, Koonin EV. 2001. The DNA-repair protein AlkB, EGL-9, and leprecan define new families of 2-oxoglutarate- and iron-dependent dioxygenases. Genome Biol. 2:research0007.1- research0007.8. doi: 10.1186/gb-2001-2-3-research0007.

Bedoya LC, Daròs J-A. 2010. Stability of Tobacco etch virus infectious clones in plasmid vectors. Virus Res. 149:234–240. doi: 10.1016/j.virusres.2010.02.004.

Bratlie MS, Drabløs F. 2005. Bioinformatic mapping of AlkB homology domains in viruses. BMC Genomics. 6:1. doi: 10.1186/1471-2164-6-1.

Canchaya C, Proux C, Fournous G, Bruttin A, Brussow H. 2003. Prophage Genomics. Microbiol. Mol. Biol. Rev. 67:238–276. doi: 10.1128/MMBR.67.2.238-276.2003.

Carrasco P, Daròs J-A, Agudelo-Romero P, Elena SF. 2007. A real-time RT-PCR assay for quantifying the fitness of tobacco etch virus in competition experiments. J. Virol. Methods. 139:181–188. doi: 10.1016/j.jviromet.2006.09.020.

Carrington JC, Haldeman R, Dolja VV, Restrepo-Hartwig MA. 1993. Internal cleavage and trans-proteolytic activities of the VPg-proteinase (NIa) of tobacco etch potyvirus in vivo. J. Virol. 67:6995–7000.

Casjens S. 2003. Prophages and bacterial genomics: what have we learned so far? Mol. Microbiol. 49:277–300. doi: 10.1046/j.1365-2958.2003.03580.x.

Cripe TP, Delos SE, Estes PA, Garcea RL. 1995. In vivo and in vitro association of hsc70 with polyomavirus capsid proteins. J. Virol. 69:7807–7813.

Díaz-Pendón JA, Ding SW. 2008. Direct and Indirect Roles of Viral Suppressors of RNA Silencing in Pathogenesis. Annu. Rev. Phytopathol. 46:303–326. doi: 10.1146/annurev.phyto.46.081407.104746.

Ding SW, Li WX, Symons RH. 1995. A novel naturally occurring hybrid gene encoded by a plant RNA virus facilitates long distance virus movement. EMBO J. 14:5762–5772.

Ding SW, Shi BJ, Li WX, Symons RH. 1996. An interspecies hybrid RNA virus is significantly more virulent than either parental virus. Proc. Natl. Acad. Sci. USA. 93:7470–7474.

Ding SW, Voinnet O. 2007. Antiviral Immunity Directed by Small RNAs. Cell. 130:413–426. doi: 10.1016/j.cell.2007.07.039.

Dolja VV, Herndon KL, Pirone TP, Carrington JC. 1993. Spontaneous mutagenesis of a plant potyvirus genome after insertion of a foreign gene. J. Virol. 67:5968–5975.

Dolja VV, Karasev A V, Koonin E V. 1994. Molecular biology and evolution of closteroviruses: sophisticated build-up of large RNA genomes. Annu. Rev. Phytopathol. 32:261–285. doi: 10.1146/annurev.py.32.090194.001401.

Dolja VV, Kreuze JF, Valkonen JPT. 2006. Comparative and functional genomics of closteroviruses. Virus Res. 117:38–51. doi: 10.1016/j.virusres.2006.02.002.

Finn RD et al. 2014. Pfam: the protein families database. Nucleic Acids Res. 42:D222–D230. doi: 10.1093/nar/gkt1223.

Gal-On A. 2000. A Point Mutation in the FRNK Motif of the Potyvirus Helper Component-Protease Gene Alters Symptom Expression in Cucurbits and Elicits Protection Against the Severe Homologous Virus. Phytopathology. 90:467–473. doi: 10.1094/PHYTO.2000.90.5.467.

Gammelgard E, Mohan M, Valkonen JPT. 2007. Potyvirus-induced gene silencing: the dynamic process of systemic silencing and silencing suppression. J. Gen. Virol. 88:2337–2346. doi: 10.1099/vir.0.82928-0.

Gilbert C et al. 2014. Population genomics supports baculoviruses as vectors of horizontal transfer of insect transposons. Nat. Commun. 5:3348. doi: 10.1038/ncomms4348.

Guo HS, Ding SW. 2002. A viral protein inhibits the long range signaling activity of the gene silencing signal. EMBO J. 21:398–407. doi: 10.1093/emboj/21.3.398.

Holmes EC. 2003. Error thresholds and the constraints to RNA virus evolution. Trends Microbiol. 11:543–546. doi: 10.1016/j.tim.2003.10.006.

Kasschau KD, Cronin S, Carrington JC. 1997. Genome Amplification and Long-Distance Movement Functions Associated with the Central Domain of Tobacco Etch Potyvirus Helper Component–Proteinase. Virology. 228:251–262. doi: 10.1006/viro.1996.8368.

Kelley WL. 1998. The J-domain family and the recruitment of chaperone power. Trends Biochem. Sci. 23:222–227.

Koboldt DC et al. 2012. VarScan 2: Somatic mutation and copy number alteration discovery in cancer by exome sequencing. Genome Res. 22:568–576. doi: 10.1101/gr.129684.111.

Kung YJ et al. 2014. Genetic analyses of the FRNK motif function of Turnip mosaic virus uncover multiple and potentially interactive pathways of cross-protection. Mol. Plant-Microbe Interact. 27:944–955. doi: 10.1094/MPMI-04-14-0116-R.

Kurowski MA, Bhagwat AS, Papaj G, Bujnicki JM. 2003. Phylogenomic identification of five new human homologs of the DNA repair enzyme AlkB. BMC Genomics. 4:48. doi: 10.1186/1471-2164-4-48.

Langmead B, Salzberg SL. 2012. Fast gapped-read alignment with Bowtie 2. Nat. Methods. 9:357–359. doi: 10.1038/nmeth.1923.

Li HW et al. 1999. Strong host resistance targeted against a viral suppressor of the plant gene silencing defence mechanism. EMBO J. 18:2683–2691. doi: 10.1093/emboj/18.10.2683.

Li H et al. 2009. The sequence alignment/map format and SAMtools. Bioinformatics. 25:2078–2079. doi: 10.1093/bioinformatics/btp352.

Lin SS, Wu HW, Jan FJ, Hou RF, Yeh SD. 2007. Modifications of the Helper Component-Protease of Zucchini yellow mosaic virus for generation of attenuated mutants for cross protection against severe infection. Phytopathology. 97:287–296. doi: 10.1094/PHYTO-97-3-0287.

Liu H et al. 2010. Widespread horizontal gene transfer from double-stranded RNA viruses to eukaryotic nuclear genomes. J. Virol. 84:11876–11887. doi: 10.1128/JVI.00955-10.

Majer E, Daròs J-A, Zwart M. 2013. Stability and fitness impact of the visually discernible Rosea1 marker in the Tobacco etch virus genome. Viruses. 5:2153–2168. doi:10.3390/v5092153.

Moreira D, Brochier-Armanet C. 2008. Giant viruses, giant chimeras: the multiple evolutionary histories of Mimivirus genes. BMC Evol. Biol. 8:12. doi: 10.1186/1471-2148-8-12.

Muniesa M, Colomer-Lluch M, Jofre J. 2013. Potential impact of environmental bacteriophages in spreading antibiotic resistance genes. Future Microbiol. 8:739–751. doi: 10.2217/fmb.13.32.

Penadés JR, Chen J, Quiles-Puchalt N, Carpena N, Novick RP. 2015. Bacteriophage-mediated spread of bacterial virulence genes. Curr. Opin. Microbiol. 23:171–178. doi: 10.1016/j.mib.2014.11.019.

Plisson C et al. 2003. Structural characterization of HC-Pro, a plant virus multifunctional protein. J. Biol. Chem. 278:23753–23761. doi: 10.1074/jbc.M302512200.

R Core Team. 2014. R: A language and environment for statistical computing. http://www.r-project.org/ (last accessed October 9, 2016).

Revers F, Le Gall O, Candresse T, Maule AJ. 1999. New advances in understanding the molecular biology of plant/potyvirus interactions. Mol. Plant-Microbe Interact. 12:367–376. doi: 10.1094/MPMI.1999.12.5.367.

Schmidt GW, Delaney SK. 2010. Stable internal reference genes for normalization of real-time RT-PCR in tobacco (Nicotiana tabacum) during development and abiotic stress. Mol. Genet. Genomics. 283:233–241. doi: 10.1007/s00438-010-0511-1.

Schmieder R, Edwards R. 2011. Quality control and preprocessing of metagenomic datasets. Bioinformatics. 27:863–864. doi: 10.1093/bioinformatics/btr026.

Sedgwick B, Bates P, Paik J, Jacobs S, Lindahl T. 2007. Repair of alkylated DNA: Recent advances. DNA Repair (Amst). 6:429–442. doi: 10.1016/j.dnarep.2006.10.005.

Shi BJ, Miller J, Symons RH, Palukaitis P. 2003. The 2b protein of cucumoviruses has a role in promoting the cell-to-cell movement of pseudorecombinant viruses. Mol. Plant-Microbe Interact. 16:261–267. doi: 10.1094/MPMI.2003.16.3.261.

Shiboleth YM et al. 2007. The conserved FRNK box in HC-Pro, a plant viral suppressor of gene silencing, is required for small RNA binding and mediates symptom development. J. Virol. 81:13135–13148. doi: 10.1128/JVI.01031-07.

Spencer DH et al. 2014. Performance of common analysis methods for detecting low-frequency single nucleotide variants in targeted next-generation sequence data. J. Mol. Diagnostics. 16:75– 88. doi: 10.1016/j.jmoldx.2013.09.003.

Stenger DC, French R, Gildow FE. 2005. Complete Deletion of Wheat Streak Mosaic Virus HC- Pro: a Null Mutant Is Viable for Systemic Infection. J. Virol. 79:12077–12080. doi: 10.1128/JVI.79.18.12077-12080.2005.

Stenger DC, Hein GL, French R. 2006. Nested deletion analysis of Wheat streak mosaic virus HC-Pro: Mapping of domains affecting polyprotein processing and eriophyid mite transmission. Virology. 350:465–474. doi: 10.1016/j.virol.2006.02.015.

Susaimuthu J, Tzanetakis IE, Gergerich RC, Martin RR. 2008. A member of a new genus in the Potyviridae infects Rubus. Virus Res. 131:145–151. doi: 10.1016/j.virusres.2007.09.001.

Torres-Barceló C, Martín S, Daròs J-A, Elena SF. 2008. From hypo- to hypersuppression: effect of amino acid substitutions on the RNA-silencing suppressor activity of the Tobacco etch potyvirus HC-Pro. Genetics. 180:1039–1049. doi: 10.1534/genetics.108.091363.

Torres-Barceló C, Daròs J-A, Elena SF. 2010. Compensatory molecular evolution of HC-Pro, an RNA-silencing suppressor from a plant RNA virus. Mol. Biol. Evol. 27:543–51. doi: 10.1093/molbev/msp272.

Urcuqui-Inchima S, Haenni a L, Bernardi F. 2001. Potyvirus proteins: a wealth of functions. Virus Res. 74:157–175.

van den Born E et al. 2008. Viral AlkB proteins repair RNA damage by oxidative demethylation. Nucleic Acids Res. 36:5451–5461. doi: 10.1093/nar/gkn519.

Varrelmann M, Maiss E, Pilot R, Palkovics L. 2007. Use of pentapeptide-insertion scanning mutagenesis for functional mapping of the plum pox virus helper component proteinase suppressor of gene silencing. J. Gen. Virol. 88:1005–1015. doi: 10.1099/vir.0.82200-0.

Verchot J, Carrington JC. 1995a.) Debilitation of plant potyvirus infectivity by P1 proteinase-inactivating mutations and restoration by second-site modifications. J. Virol. 69:1582–1590.

Verchot J, Carrington JC. 1995b.) Evidence that the potyvirus P1 proteinase functions in trans as an accessory factor for genome amplification. J. Virol. 69:3668–3674.

Willemsen A, Zwart MP, et al. 2016a.) Multiple Barriers to the Evolution of Alternative Gene Orders in a Positive-Strand RNA Virus. Genetics. 202:1503–1521. doi: 10.1534/genetics.115.185017.

Willemsen A, Zwart MP, Higueras P, Sardanyés J, Elena SF. 2016b. Predicting the stability of homologous gene duplications in a plant RNA virus. Genome Biol. Evol. In Press. doi: 10.1093/gbe/evw219.

Wu HW, Lin S-S, Chen KC, Yeh S-D, Chua N-H. 2010. Discriminating mutations of HC-Pro of zucchini yellow mosaic virus with differential effects on small RNA pathways involved in viral pathogenicity and symptom development. Mol. Plant-Microbe Interact. 23:17–28. doi: 10.1094/MPMI-23-1-0017.

Young BA, Hein GL, French R, Stenger DC. 2007. Substitution of conserved cysteine residues in wheat streak mosaic virus HC-Pro abolishes virus transmission by the wheat curl mite. Arch. Virol. 152:2107–2111. doi: 10.1007/s00705-007-1034-x.

Zwart MP, Willemsen A, Daròs J-A, Elena SF. 2014. Experimental evolution of pseudogenization and gene loss in a plant RNA virus. Mol. Biol. Evol. 31:121–134. doi: 10.1093/molbev/mst175.

